# High-throughput CRISPRi phenotyping in *Streptococcus pneumoniae* identifies new essential genes involved in cell wall synthesis and competence development

**DOI:** 10.1101/088336

**Authors:** Xue Liu, Clement Gallay, Morten Kjos, Arnau Domenech, Jelle Slager, Sebastiaan P. van Kessel, Kèvin Knoops, Robin A. Sorg, Jing-Ren Zhang, Jan-Willem Veening

## Abstract

Genome-wide screens have discovered a large set of essential genes in the opportunistic human pathogen *Streptococcus pneumoniae*. However, the functions of many essential genes are still unknown, hampering vaccine development and drug discovery. Based on results from transposon-sequencing (Tn-Seq), we refined the list of essential genes in *S. pneumoniae* serotype 2 strain D39. Next, we created a knockdown library targeting 348 potentially essential genes by CRISPR interference (CRISPRi) and show a growth phenotype for 254 of them (73%). Using high-content microscopy screening, we searched for essential genes of unknown function with clear phenotypes in cell morphology upon CRISPRi-based depletion. We show that SPD1416 and SPD1417 (renamed to MurT and GatD, respectively) are essential for peptidoglycan synthesis, and that SPD1198 and SPD1197 (renamed to TarP and TarQ, respectively) are responsible for the polymerization of teichoic acid (TA) precursors. This knowledge enabled us to reconstruct the unique pneumococcal TA biosynthetic pathway. CRISPRi was also employed to unravel the role of the essential Clp-proteolytic system in regulation of competence development and we show that ClpX is the essential ATPase responsible for ClpP-dependent repression of competence. The CRISPRi library provides a valuable tool for characterization of pneumococcal genes and pathways and revealed several promising antibiotic targets.

## Introduction

*Streptococcus pneumoniae* (pneumococcus) is a major cause of community-acquired pneumonia, meningitis and acute otitis media and, despite the introduction of several vaccines, remains one of the leading bacterial causes of mortality worldwide (Prina et al, 2015). The main antibiotics used to treat pneumococcal infections belong to the beta-lactam class, such as amino-penicillins (amoxicillin, ampicillin) and cephalosporines (cefotaxime). These antibiotics target the penicillin binding proteins (PBPs), which are responsible for the synthesis of peptidoglycan (PG) that plays a role in the maintenance of cell integrity, cell division and anchoring of surface proteins (Kocaoglu et al, 2015; Sham et al, 2012). The pneumococcal cell wall furthermore consists of teichoic acids (TA), which are anionic glycopolymers that are either anchored to the membrane (lipo TA) or covalently attached to PG (wall TA) and are essential for maintaining cell shape (Brown et al, 2013; Massidda et al, 2013). Unfortunately, resistance to most beta-lactam antibiotics remains alarmingly high. For example, penicillin non-susceptible pneumococcal strains colonizing the nasopharynx of children remains above 40 % in the United States (Kaur et al, 2016), despite the effect of the pneumococcal conjugate vaccines. The introduction of several conjugate vaccines during the last two decades, effectively reduced nearly all the strains expressing capsules targeted by the vaccines, reducing the number of resistant infections; nevertheless, these vaccines target up to 13 of 96 described serotypes, and clones expressing new serotypes have rapidly emerged and spread worldwide (Kim et al., 2016). In fact, the frequency of pneumococcal colonization remained stable in spite of the introduction of vaccines (Kaur et al, 2016). Furthermore, multidrug resistance in *S. pneumoniae* is prevalent and antibiotic resistance determinants and virulence factors can readily transfer between strains via competence-dependent horizontal gene transfer (Chewapreecha et al, 2014; Johnston et al, 2014; Kim et al, 2016). For these reasons, it is crucial to understand how competence is regulated and to identify and characterize new essential genes and pathways. Interestingly, not all proteins within the pneumococcal PG and TA biosynthesis pathways are known (Massidda et al, 2013), leaving room for discovery of new potential antibiotic targets. For instance, not all enzymes in the biosynthetic route to lipid II, the precursor of PG, are known and annotated in *S. pneumoniae*. The pneumococcal TA biosynthetic pathway is even more enigmatic and it is unknown which genes code for the enzymes responsible for polymerizing TA precursors (Denapaite et al, 2012).

Several studies using targeted gene knockout and depletion/overexpression techniques as well as transposon sequencing (Tn-Seq), have aimed to identify the core pneumococcal genome (Mobegi et al, 2014; Song et al, 2005; Thanassi et al, 2002; van Opijnen et al, 2009; van Opijnen & Camilli, 2012; Verhagen et al, 2014; Zomer et al, 2012). These genome-wide studies revealed a core genome of around 400 genes essential for growth either *in vitro* or *in vivo*. Most of the essential pneumococcal genes can be assigned to a functional category on basis of sequence homology or experimental evidence. However, approximately one third of the essential genes belong to the category of ‘function unknown’ or ‘hypothetical’ and it is likely that several unknown cell wall synthesis genes, such as the TA polymerase, are present within this category.

To facilitate the high-throughput study of essential genes in *S. pneumoniae* on a genome-wide scale, we established CRISPRi (clustered regularly interspaced short palindromic repeats interference) for this organism. CRISPRi is based on expression of a nuclease-inactive *Streptococcus pyogenes* Cas9 (dCas9), which, together with expression of a single-guide RNA (sgRNA) targets the gene of interest (Peters et al, 2016; Qi et al, 2013). When targeting the non-template strand of a gene by complementary base-pairing of the sgRNA with the target DNA, the dCas9-sgRNA-DNA complex acts as a roadblock for RNA polymerase (RNAP) and thereby represses transcription of the target genes (Peters et al, 2016; Qi et al, 2013) (Fig. 1A). Note that *S. pneumoniae* does not contain an endogenous CRISPR/Cas system, likely because it interferes with natural transformation and thereby lateral gene transfer that is crucial for pneumococcal host adaptation (Bikard et al, 2012).

**Figure 1.**
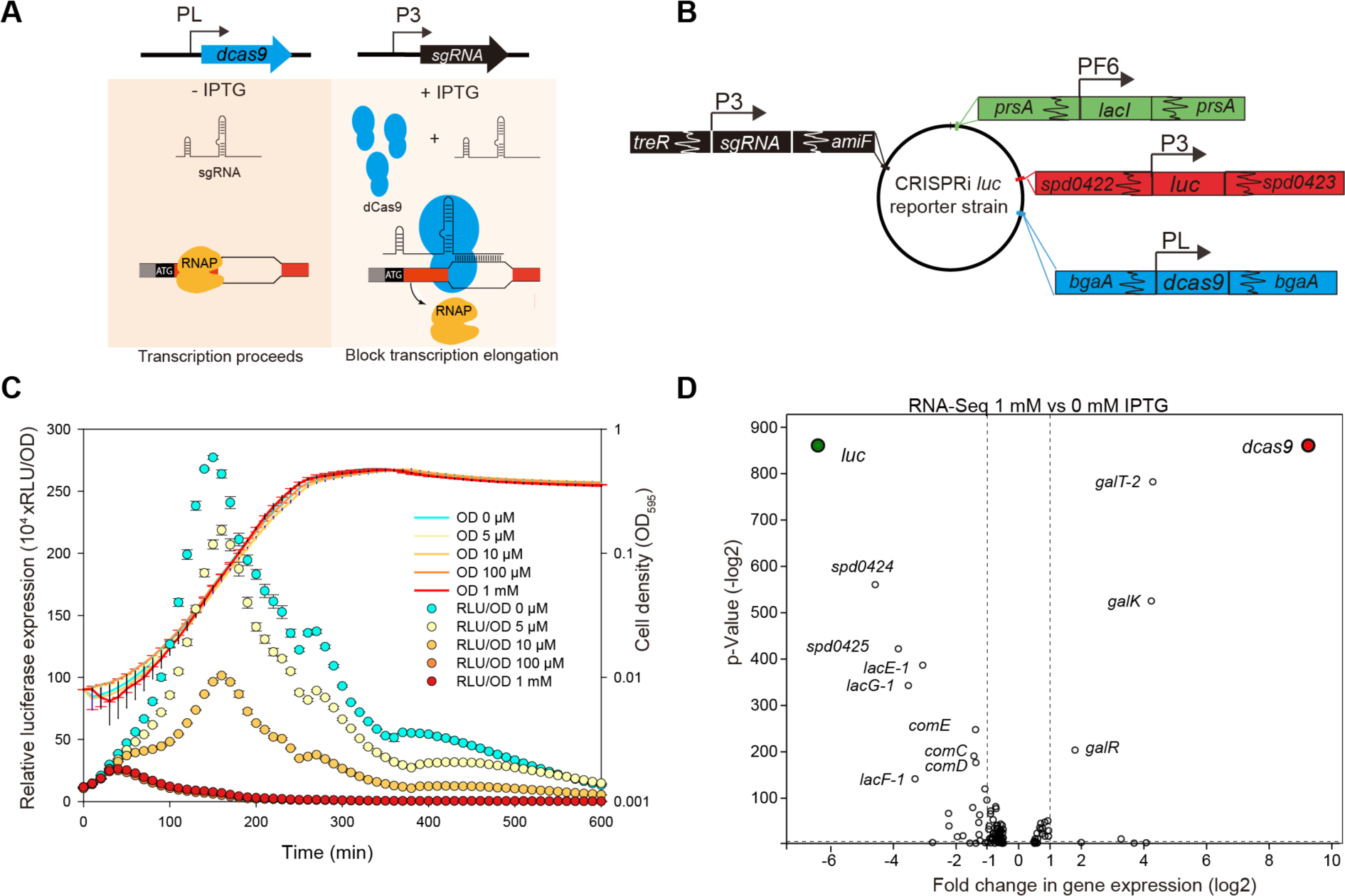
An IPTG-inducible CRISPRi system for tunable repression of gene expression in *S. pneumoniae*. **A**. *dcas9* and sgRNA sequences were chromosomally integrated at two different loci and expression was driven by an IPTG-inducible promoter (PL) and a constitutive promoter (P3), respectively. With addition of IPTG, dCas9 is expressed and guided to the target site by constitutively expressed sgRNA. Binding of dCas9 to the 5’ end of the coding sequence of its target gene blocks transcription elongation. In the absence of IPTG, expression of dCas9 is tightly repressed, and transcription of the target gene can proceed smoothly. **B**. Genetic map of CRISPRi *luc* reporter strain XL28. To allow IPTG-inducible expression, the *lacI* gene, driven by the constitutive PF6 promoter, was inserted at the non-essential *prsA* locus; *luc,* encoding firefly luciferase, driven by the constitutive P3 promoter was inserted into the intergenic sequence between gene loci *spd0422* and *spd0423*; *dcas9* driven by the IPTG-inducible PL promoter was inserted into the *bgaA* locus; sgRNA-*luc* driven by the constitutive P3 promoter was inserted into CEP locus (between *treR* and *amiF*). **C**. The CRISPRi system tested in the *luc* reporter strain XL28. Expression of *dcas9* was induced by addition of different concentrations of IPTG. Cell density (OD595) and luciferase activity (shown as RLU/OD) of the bacterial culture were measured every 10 minutes. The value represents averages of three replicates with SEM. **D**. RNA-seq confirms the specificity of the CRISPRi system in *S. pneumoniae*. RNA sequencing was performed on the *luc* reporter strain XL28 (panel B) with or without 1 mM IPTG. The *dcas9* and *luc* genes are highlighted. Data were analyzed with T-REx and plotted as a Volcano plot. *p*-Value equals 0.05 is represented by the horizontal dotted line. Two vertical dotted lines mark the 2 fold changes.

Using Tn-Seq and CRISPRi, we refined the list of genes that are either essential for viability or for fitness in the commonly used *S. pneumoniae* strain D39 (Avery et al, 1944). To identify new genes involved in pneumococcal cell envelope homeostasis, we screened for essential genes of unknown function with a clear morphological defect upon CRISPRi-based depletion. This identified SPD1416 and SPD1417 as essential peptidoglycan synthesis proteins (renamed to MurT and GatD, respectively) and SPD1198 and SPD1197 as essential proteins responsible for precursor polymerization in TA biosynthesis (hereafter called TarP and TarQ, respectively). Finally, we demonstrate the use of CRISPRi to unravel gene regulatory networks and showed that ClpX is the ATPase subunit that acts together with the ClpP protease as a repressor for competence development.

## Results and Discussion

### Identification of potentially essential genes in *S. pneumoniae*strain D39

While several previous studies have identified many pneumococcal genes that are likely to be essential, the precise contribution to pneumococcal biology has remained to be defined for most of these genes. Here, we aim to characterize the functions of these proteins in the commonly used *S. pneumoniae* serotype 2 strain D39 by the CRISPRi approach. Therefore, we performed Tn-Seq on *S. pneumoniae* D39 grown in C+Y medium at 37ºC, our standard laboratory condition (see Methods). By combining our D39 data with the results from two Tn-Seq studies performed in serotype 4 strain TIGR4 (van Opijnen et al, 2009; van Opijnen & Camilli, 2012), we calculated the essentiality score for each gene and selected 391 potentially essential genes (Supplementary Table 1), to be included in our pneumococcal essential gene CRISPRi library (see below).

### CRISPRi enables tunable repression of gene transcription in *S. pneumoniae*

To develop the CRISPR interference system, we first engineered the commonly used LacI-based isopropyl β-D-1-thiogalactopyranoside (IPTG)-inducible system for *S. pneumoniae* (see Methods). *dcas9* was placed under control of this new IPTG-inducible promoter, named PL, and was integrated into the chromosome via double crossover (Fig. 1A-B). To confirm the reliability of the CRISPRi system, we tested it in a reporter strain expressing firefly luciferase (*luc*), in which a sgRNA targeting *luc* was placed under the constitutive P3 promoter (Sorg et al, 2015) and integrated at a non-essential locus (Fig. 1B). To obtain high efficiency of transcriptional repression, we used the optimized sgRNA sequence as reported previously (Chen et al, 2013) (Fig. S1A).

Induction of dCas9 with 1 mM IPTG resulted in quick reduction of luciferase activity; approximately 30-fold repression of luciferase expression was obtained within 2 hr without substantial impact on bacterial growth (Fig. 1C). Furthermore, the level of repression was tunable by using different concentrations of IPTG (Fig. 1C). By comparing strains with or without sgRNA*luc*, we found that repression in our CRISPRi system is stringently dependent on expression of both dCas9 and the sgRNA (Fig. S1B). To test the precision of CRISPRi in *S. pneumoniae*, we determined the transcriptome of the sgRNA*luc* strain (Strain XL28) by RNA-Seq in the presence or absence of IPTG. The data was analyzed using Rockhopper (McClure et al, 2013) and T-REx (de Jong et al, 2015). The RNA-Seq data showed that expression of dCas9 was stringently repressed by LacI without IPTG, and was upregulated ~600 fold upon addition of 1 mM IPTG after 2.5 hr. Upon dCas9 induction, the *luc* gene was significantly repressed (~ 84 fold) (Fig. 1D). Our RNA-seq data showed that the genes (*spd0424*, *spd0425*, *lacE-1*, *lacG-1*, *lacF-1*) that are downstream of *luc,* which was driven by a strong constitutive promoter without terminator, were significantly repressed as well, confirming the reported polar effect of CRISPRi (Qi et al, 2013). Interestingly, by comparing the transcriptome of strain with (XL28) or without (XL29) sgRNA*luc* cultivated in medium with 1 mM IPTG, we found that *galT-2*, *galK* and *galR* were specifically induced by IPTG, and fluctuation of the competence system was also observed which is known to be highly noisy (Aprianto et al, 2016; Prudhomme et al, 2016) (Supplementary Table 2). Taken together, the IPTG-inducible CRISPRi system is highly specific.

### Construction and growth analysis of the CRISPRi library

We next used the CRISPRi system to construct an expression knock-down library of pneumococcal essential genes. A sgRNA to each of the 391 potentially essential genes was designed as described previously (Larson et al, 2013) (Supplementary Table 3). Note that CRISPRi also inhibits expression of co-transcribed downstream genes (Fig. 1D) (Qi et al, 2013). Thus, some of the genes may be targeted multiple times (in case of more than one essential gene within the operon). We used an infusion cloning technique to construct the library as this was more efficient than the previously reported PCR-based sgRNA construction method (see SI text) (Figs 2A and 2B). All sgRNA strains were sequence verified, and we considered them genetically functional when the sgRNA did not contain more than 1 mismatch to the designed sgRNA and no mismatches in the first 14-nt prior to the PAM. Using this approach, after a single round of cloning and sequencing, we successfully constructed 348 unique sgRNA strains (see Methods). Note that we are still in the process of constructing the remaining 43 sgRNA strains.

To examine the effects of CRISPRi-based gene silencing, growth was assayed both in the presence and absence of 1 mM IPTG for 18 hours in real-time by microtiterplate assays. Two types of growth phenotypes were defined and identified: a growth defect and increased lysis (Figs S2B and S2C). As shown in Fig 2C, CRISPRi-based repression of transcription of 230 unique genes led to a growth defect, 48 genes showed increased lysis, including 24 that demonstrated both a growth defect and increased lysis, and 94 genes showed no defect (see supplementary Table 1). In total, 254 out of 348 target genes (about 73%) repressed by CRISPRi showed growth phenotypes. Comparing the optical densities between the uninduced and induced cells at the time point at which uninduced cells reached an OD595 of 0.1, 174 genes repressed by CRISPRi displayed a more than 4-fold growth defect, and 254 genes showed a more than 2-fold growth defect (Fig. 2D).

**Figure 2.**
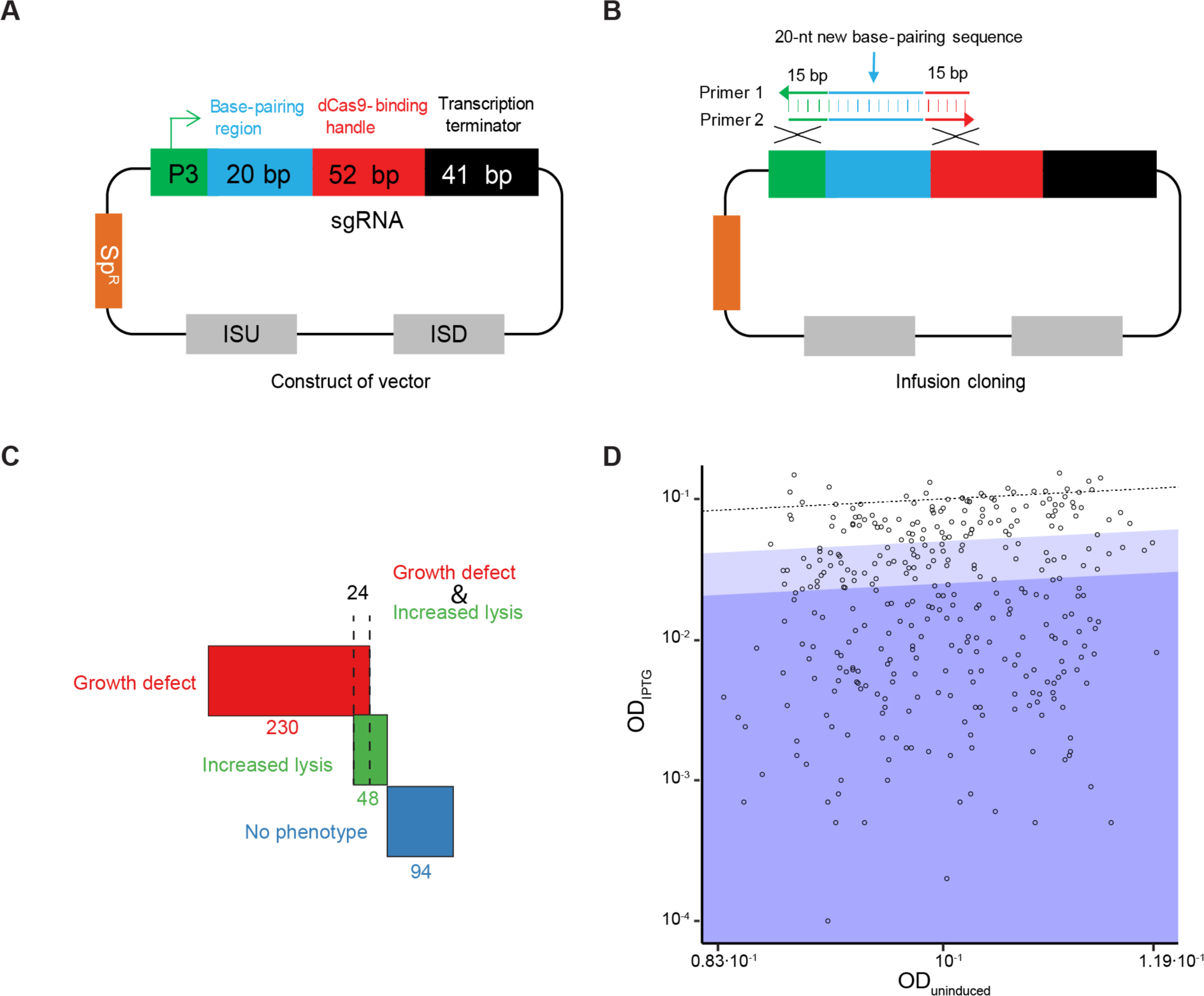
Construction and growth analysis of the CRISPRi library. **A**. The plasmid map of the sgRNA vector. The sgRNA expression vector is a *S. pneumoniae* integration vector. It contains a constitutive P3 promoter, a spectinomycin-selectable marker (SpR), two homology sequences (ISU and ISD) for double crossover integration, and the sgRNA sequence. The sgRNA chimera contains a base-pairing region (blue), dCas9-binding handle (red) and the *S. pyogenes* transcription terminator (black). **B**. Schematic of the infusion method used for sgRNA-cloning. Two gene-specific primers were designed for each cloning. Primer 1 and primer 2 are complementary, and they contain 15-bp homology sequences to the adjoining region, flanking the 20-bp base-pairing region of the sgRNA encoding sequence of the vector. **C and D**. Growth defect analysis of the whole library. **C**. Classification of the 348 genes targeted by the CRISPRi library according to growth analysis. Criteria for determination of a growth defect and increased lysis is demonstrated in Figs. S2B and S2C. **D**. OD595 of IPTG-induced cells is plotted against the OD595 of uninduced cells for a representative dataset from each of the 348 genes under study. The timepoint at which uninduced cells have an Optical Density (595nm) closest to 0.1 was selected for plotting. Data points in the dark blue area (174/348 strains) correspond to genes displaying a strong growth defect (more than 4-fold); points in the light blue area induced cells display a moderate growth defect of 2-to 4-fold (71/348 strains). The same type of analysis was performed on 36 negative control strains, shown in Fig. S2D.

### Phenotyping important pneumococcal genes by combined CRISPRi and high-content microscopy

To test whether CRISPRi was able to place genes in a functional category and thereby allow us to identify previously uncharacterized genes with a function in cell envelope homeostasis, we first analyzed the effects of CRISPRi-based repression on cell morphology using 69 genes that have been reported to be essential or crucial for normal pneumococcal growth. These genes have been associated with capsule synthesis (3 genes), transcription (4 genes), cell division (6 genes), translation (7 genes), teichoic acid biosynthesis (10 genes), cell membrane synthesis (11 genes), chromosome biology (14 genes), and peptidoglycan synthesis (14 genes) (Fig. 3, Table 1, Figs S3-S10). This showed a good correlation between reported gene function and observed phenotype (SI text). For instance, repression of transcription of genes involved in chromosome biology caused, as expected, appearance of anucleate cells or cells with aberrant chromosomes (Fig. S3). Importantly, repression of transcription of genes known to be involved in cell envelope homeostasis such as *ftsZ*, *ftsL*, *ftsW*, *rodA*, *pbp2X*, *glmU*, *murC, murF*, *tarI*, *tarJ*, *licA*, *licB*, *licC and licD3*, caused severe changes in cell shape and size, including heterogeneous cell size, ballooning cells, defective septa, short cells, round cells, cells in chains and cells demonstrating a coccus-to-rod transition (Figs S7, S9 and S10).

**Figure 3.**
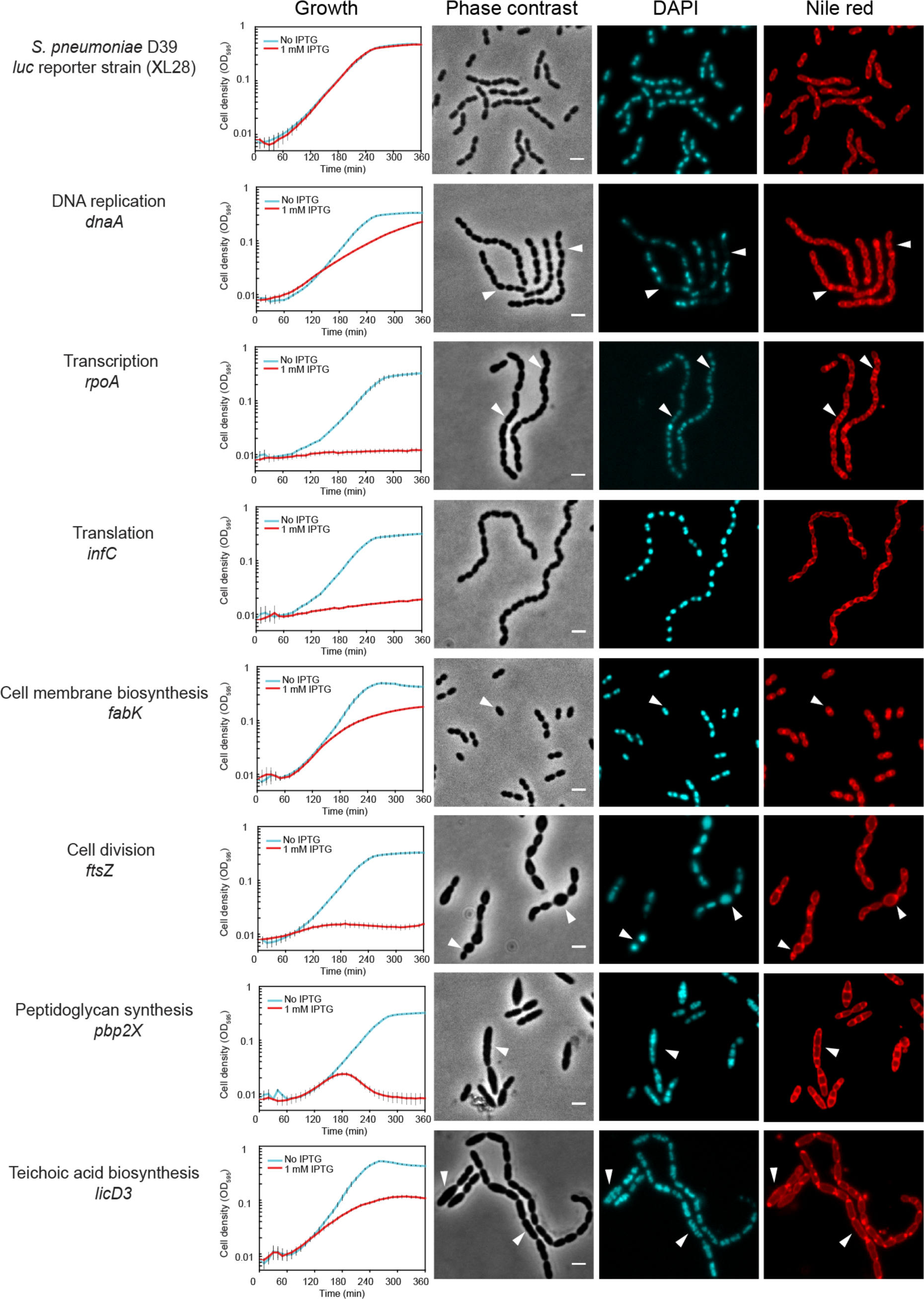
Growth profiles and morphological changes of CRISPRi strains with sgRNA targeting genes of different functional pathways. Growth of *S. pneumoniae* strains was performed in C+Y medium with (red) or without (cyan) 1 mM IPTG. Morphological changes were examined with fluorescence microscopy and representative micrographs are shown. Phase contrast, DAPI staining DNA and nile red staining membrane are displayed. Scale bar = 2 ¼m. White arrows point to the typical morphological changes. *S. pneumoniae* D39 reporter strain XL28 expresses firefly luciferase (*luc)* from a constitutive promoter and contains sgRNA targeting the *luc* gene.

**Table 1.**
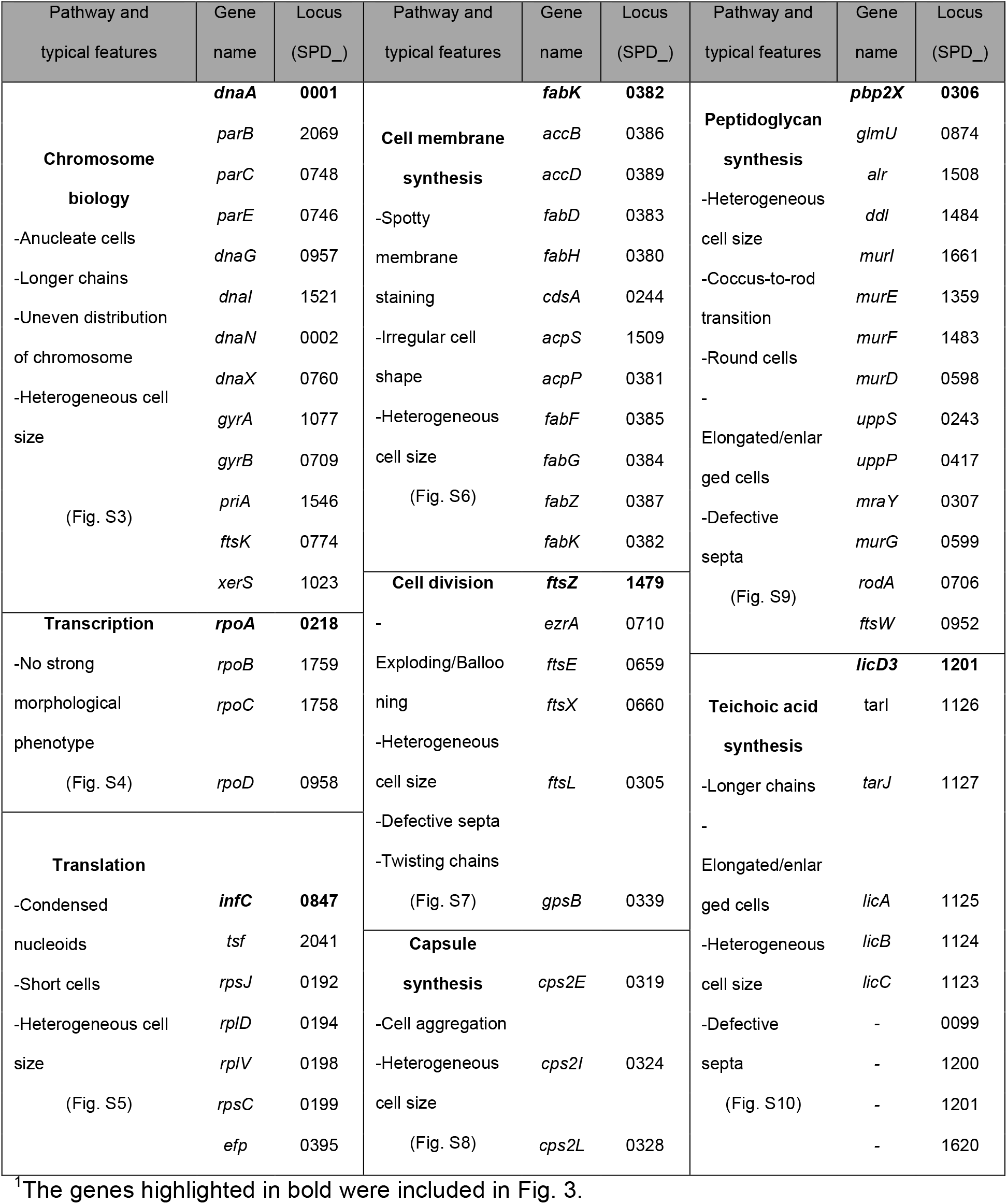
Cellular pathways selected for CRISPRi phenotyping^1^.

Using the same approach, we could also functionally verify several genes that were studied before but had not been annotated in the D39 genome. These included *pcsB* (*spd2043*), *vicR* (*spd1085*), *divIC* (*spd0008*) *and rafX* (*spd1672*). Transcriptional repression led to significant growth defects and cellular changes in shape and size (SI text and Fig. S12). Similarly, repression of *spd1405*, *spd1522* and *spd0827* led to significant growth defects and generation of anucleate cells. Our data revealed that these “hypothetical” genes represent the functional orthologs of *dnaB* (*spd1405*), *dnaD* (*spd1522*) and *yabA* (*spd0827*) in *S. pneumoniae* (SI text and Fig. S13).

### SPD1416 and SPD1417 are involved in peptidoglycan precursor synthesis

Using high-throughput fluorescence microscopy, we phenotyped a subset of the 348 CRISPRi knockdowns that target genes of unknown function. CRISPRi strains with sgRNA targeting hypothetical genes *spd1416* or *spd1417* showed significant growth retardation and morphological abnormality, such as heterogeneous cell size and elongated and enlarged cells with multiple incomplete septa (Fig. S11). These manifestations mirrored what we observed upon inhibiting the expression of genes known to be involved in peptidoglycan (PG) synthesis (Fig. S9). Consistent with the essentiality of these two genes as suggested by Tn-Seq, we were unable to obtain deletion mutants of *spd1416* or *spd1417* after multiple attempts. To confirm that these genes are essential for pneumococcal growth, we constructed merodiploid strains of *spd1416* and *spd1417* by inserting a second copy of each gene fused to *gfp* (encoding a monomeric superfolder GFP) at their N-terminus. These *gfp*-fusions were integrated at the ectopic *bgaA* locus under the control of the zinc-inducible promoter, PZn. In the presence of Zn^2+^, we were able to delete the native *spd1416* or *spd1417* gene by allelic replacement. While both the *spd1416* and *spd1417* mutants behaved normally in the presence of Zn^2+^, severe growth retardation was observed in the absence of Zn^2+^ (Fig. 4A). Interestingly, we did not obtain erythromycin resistant colonies when transforming in the PZn-*spd1417*-*gfp* genetic background, indicating that the C-terminal GFP fusion is not functional. Together, these lines of evidence demonstrate that both *spd1416* and *spd1417* are essential genes.

**Figure 4.**
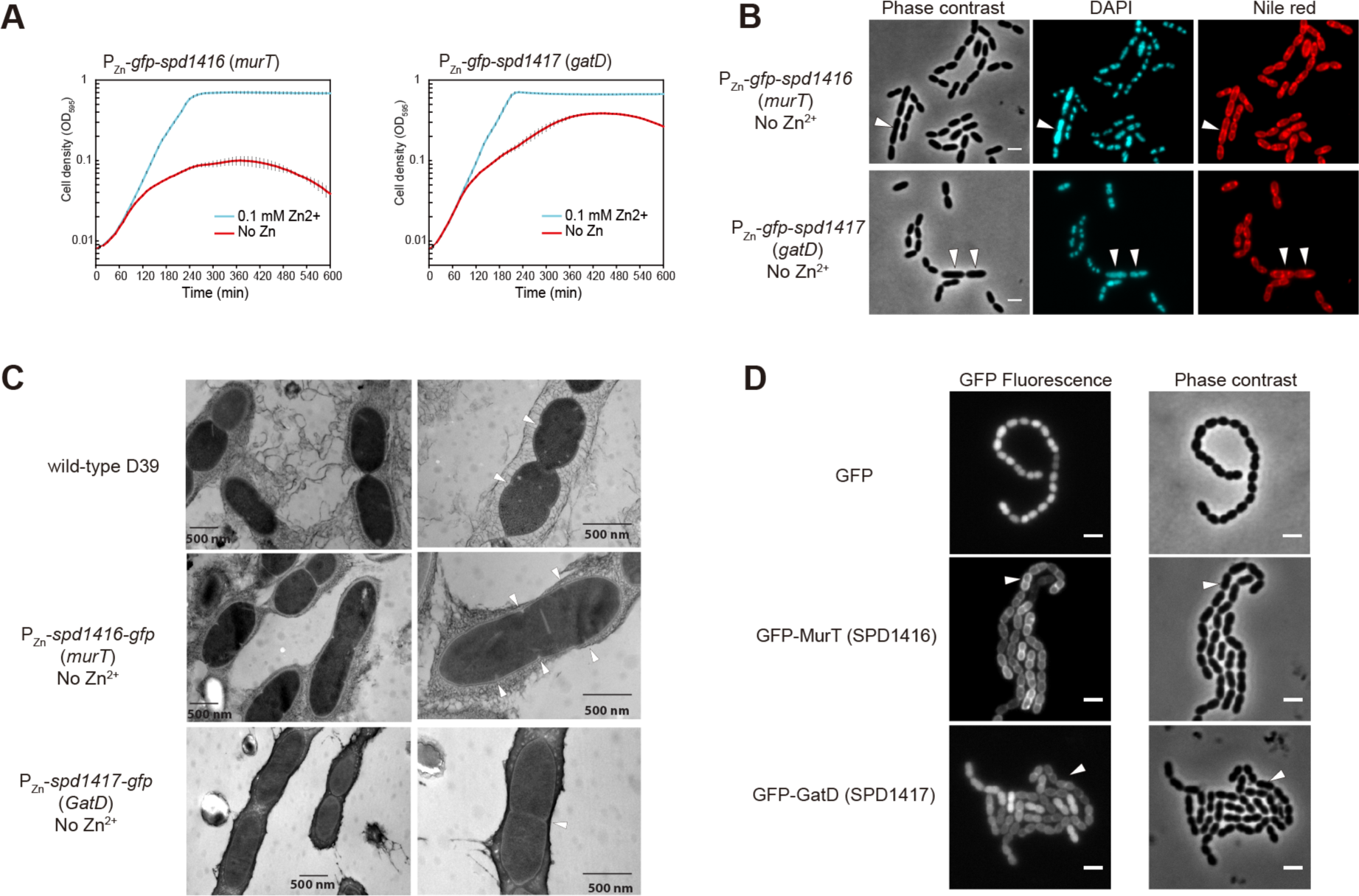
Identification of peptidoglycan synthesis genes *spd1416* (*murT*) and *spd1417* (*gatD*). **A**. Growth curves of depletion strains P*Zn*-*gfp*-*spd1416* (*murT*) and P*Zn-gfp*-*spd1417* (*gatD*), in C+Y medium with (cyan) or without (red) 0.1 mM Zn^2+^. **B**. Microscopy of cells from panel A after incubating in C+Y medium without Zn^2+^ for 2.5 hours. Representative micrographs of phase contrast, DAPI, and nile red are shown. Scale bar = 2 ¼m. White arrows point to elongated/enlarged cells. **C**. Electron micrographs of the same samples as in panel B and wild type *S. pneumoniae* D39 as reference. Note that depletion of *spd1416* or *spd1417* resulted in elongated cells. Septa are pointed with white arrows. **D**. Localization of GFP-MurT and GFP-GatD. Micrographs of GFP signal (upper panel) and phase contrast (lower panel) are shown. Scale bar = 2 ¼m. Cells pointed by white arrows show typical membrane localization. *S. pneumoniae* D39 with free GFP with cytoplasmic localization was included as reference.

Morphological analysis by light microscopy of bacterial cells upon depletion of *gfp-spd1416* or *gfp-spd1417* confirmed the morphological changes as observed in the CRISPRi knockdowns (Fig. 4B). The *gfp-spd1416* or *gfp-spd1417* cells were further analyzed using freeze-substitution electron microscopy (Fig. 4C). This showed the presence of elongated cells and the frequent formation of multiple septa per cell, in contrast to wild type D39 cells which showed the typical diplococcal shape. Note that the mild sample preparation used in our freeze-substitution EM protocol also preserved the capsule, which can be readily lost during traditional EM sample preparation (Hammerschmidt et al, 2005) (Fig. 4C). BlastP analysis shows that SPD1416 contains a Mur-ligase domain with 36% sequence identity with MurT of *Staphylococcus aureus,* whereas SPD1417 possesses a glutamine amidotransferase domain with 40% sequence identity with GatD of *S. aureus*. MurT and GatD, two recently identified proteins involved in staphylococcal cell wall synthesis (Figueiredo et al, 2012; Munch et al, 2012), form a complex to perform the amidation of the D-glutamic acid in the stem peptide of PG. It was previously reported that recombinant MurT/GatD of *S. pneumoniae* R6, purified from *Escherichia coli*, indeed can amidate glutamate lipid II into iso-glutamine lipid II *in vitro* (Zapun et al, 2013). Therefore, we named *spd1416* to *murT* and *spd1417* to *gatD*. It is interesting to note that while MurT or GatD depletion strains in *S. aureus* showed reduced growth, cells exhibited normal cell morphologies (Figueiredo et al, 2012), in contrast to the strong morphological defects observed in *S. pneumoniae* D39.

MurT and GatD contain no membrane domain or signal peptide, and are thus predicted to be cytoplasmic proteins. However, fluorescence microcopy of the N-terminal GFP fused to MurT or GatD showed that they are partially membrane localized (Fig. 4D). Ingel fluorescence imaging showed that GFP-MurT and GFP-GatD were correctly expressed without any detectable proteolytic cleavage (Fig. S14). Since *in vitro* assays demonstrated that glutamate lipid II, which is anchored to the membrane by the bactoprenol hydrocarbon chain of lipid II, is a substrate of the MurT/GatD amidotransferase complex, it is reasonable to assume that membrane localization of MurT or GatD is caused by recruitment to the membrane-bound substrate. Indeed, amidation of the glutamic acid at position 2 of the peptide chain most likely occurs after formation of lipid-linked PG precursors (Rajagopal & Walker, 2016).

### CRISPRi revealed novel pneumococcal genes involved in teichoic acid biosynthesis

CRISPRi-based repression of hypothetical essential genes *spd1197* and *spd1198* led to significant growth defects and microscopy revealed chained cells with abnormal shape and size (Fig. S11). Some of the cells were elongated and enlarged. These phenotypes are consistent with the typical morphological changes caused by repression of genes in teichoic acid (TA) biosynthesis (Fig. S10). In accordance with this, analysis of the genetic context of *spd1197* and *spd1198* showed that they are in the *lic3* region, which was predicted to be a pneumococcal TA gene cluster (Denapaite et al, 2012; Kharat et al, 2008). We were unable to replace these genes with an erythromycin resistance marker, confirming their essentiality. Similar to the approach described above, we generated Zn^2+^-inducible C-terminal GFP fusions to SPD1197 and SPD1198, integrated these ectopically at the *bgaA* locus and then successfully deleted the native *spd1197* or *spd1198* genes in the presence of Zn^2+^. Plate reader assays showed strong growth impairment in the absence of Zn^2+^ (Fig. 5A). In line with the phenotypes of the CRISPRi screen, the zinc-depletion strains showed similar morphological defects, cells in chains and elongated or enlarged cell shape and size (Fig. 5B). EM analysis of depleted cells also revealed uneven distribution of multiple septa within a single cell, increased extracellular material and a rough cell surface (Fig. 5C).

**Figure 5.**
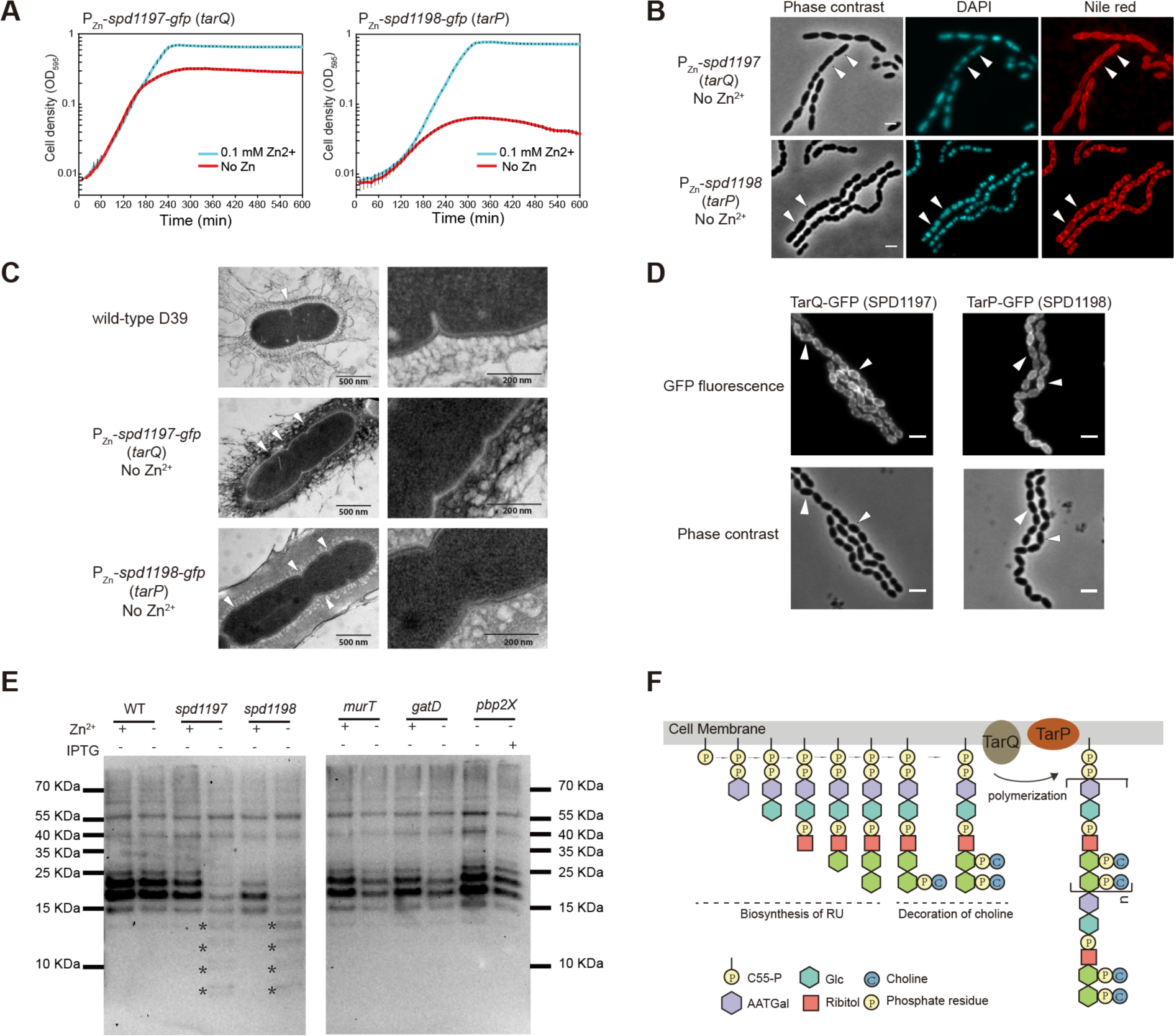
Newly identified genes of the teichoic acid biosynthesis pathway: *spd1198* (*tarP*) and *spd1197* (*tarQ*) are involved in precursor polymerization. **A**. Growth curves of depletion strains PZn-*spd1197-gfp* (*tarQ*) and PZn*-spd1198-gfp* (*tagP*) in C+Y medium with (cyan) or without (red) 0.1 mM Zn^2+^. **B**. Microscopy of cells from panel A after incubation in C+Y medium without Zn^2+^ for 2.5 hours. Representative micrographs are shown. Scale bar = 2 ¼m. White arrows point to elongated/enlarged cells. Note that depletion of *spd1197* or *spd1198* leads to chains of cells. **C**. Electron micrographs of the same samples as in panel B with wild type *S. pneumoniae* D39 as reference. Arrowheads point to the septa of cells. **D**. Localization of TarQ-GFP, TarP-GFP, with C-terminal fused monomeric GFP. GFP signal (upper panel) and phase contrast (lower panel) are shown. Scale bar = 2 ¼m. White arrows point to the cells showing typical membrane localization. **E**. Western blotting to detect phosphorylcholine-containing molecules of *S. pneumoniae*. Whole cell lysates were separated with SDS-PAGE, and phosphorylcholine containing molecules were detected by phosphorylcholine antibody TEPC-15. Smaller bands caused by depletion of *spd1197* or *spd1198* are indicated by asterisks. Note that for *spd1197*, *spd1198*, *murT*, and *gatD*, Zn^2+^ inducible strains were used, and for *pbp2X*, CRISPRi strains were used. **F**. Model for TarP/TarQ function in precursor polymerization of the teichoic acid biosynthesis pathway in *S. pneumoniae*. Steps of biosynthesis of repeat units (RU), decoration of RU with choline and polymerization of the precursor are shown.

SPD1198 contains 11 predicted transmembrane helices while SPD1197 has 2 predicted TM segments with a C-terminal extracytoplasmic tail. In-gel fluorescence showed that SPD1197-GFP was mainly produced as a full-length product. The SPD1198-GFP fusion, however, showed clear signs of protein degradation (Fig. S14). Nevertheless, we performed fluorescence microscopy to determine their localizations. In agreement with the prediction, SPD1197-GFP and SPD1198-GFP are clearly localized to the membrane (Fig. 5D).

Phosphorylcholine is an essential component of pneumococcal TA and for this reason a phosphorylcholine antibody is frequently used to detect *S. pneumoniae* TA (Vollmer & Tomasz, 2001; Wu et al, 2014). To explore whether SPD1197 and SPD1198 indeed play a role in TA synthesis, we performed Western blotting to detect phosphorylcholine-decorated TA using whole cell lysates (Fig. 5E). Cells of strains PZn-*spd1197-gfp* and PZn -*spd1198-gfp* were grown in the presence or absence of 0.1 mM Zn^2+^. As controls, we depleted expression of three genes involved in PG synthesis (*murT*, *gatD*, and *pbp2x*). As shown in Fig. 5E, Zn^2+^ did not influence TA synthesis of the *S. pneumoniae* D39 wild-type (WT) strain, and the four main TA bands are clearly visible, migrating in the range between 15 and 25 kDa consistent with previous reports (Wu et al, 2014). In contrast, cells depleted for SPD1197 or SPD1198 displayed a different pattern and the 4 main bands around 15 kDa and 25 kDa were missing or much weaker, while multiple bands with a size smaller than 15 kDa appeared. TA of *S. pneumoniae*, including wall teichoic acid (WTA) and membrane-anchored lipoteichoic acid (LTA), are polymers with identical repeating units (RU). Addition of one RU can lead to about a 1.3 kDa increase in molecular weight (Gisch et al, 2013). Interestingly, the weight interval between the extra smaller bands from bacterial cells with depleted SPD1197 or SPD1198 seemed to match the molecular weight of the RU, suggesting that SPD1197 and SPD1198 play a role in TA precursor polymerization. Although repression of the genes associated with peptidoglycan synthesis (*murT*, *gatD* and *pbp2x*) made the 4 main TA bands weaker, the pattern of the TA bands was not changed. Likely, the reduction of the TA of these three strains is due to the reduction of peptidoglycan, which constitutes the anchor for wall TA.

The TA chains of *S. pneumoniae* are thought to be polymerized before they are transported to the outside of membrane, and so far it is not known which protein(s) function(s) as TA polymerase (Denapaite et al, 2012). In line with SPD1198 being the TA polymerase, homology analysis shows that it contains a predicted polymerase domain. The large cytoplasmic part of SPD1197 may aid in the assembly of the TA biosynthetic machinery by protein-protein interactions (Denapaite et al, 2012). Together, we here show that SPD1197 and SPD1198 are essential for growth and we suggest that they are responsible for polymerization of TA chains (Fig. 5F). Consistent with the nomenclature used for genes involved in TA biosynthesis, we named *spd1198 tarP* (for teichoic <u>a</u>cid ribitol <u>p</u>olymerase) and *spd1197 tarQ* (in operon with *tarP,* sequential alphabetical order). Whether TarP and TarQ interact and function as a complex remains to be determined.

### The essential ATPase ClpX and the protease ClpP repress competence development

We wondered whether we could also employ CRISPRi to probe gene-regulatory networks in which essential genes play a role. An important pathway in *S. pneumoniae* is development of competence for genetic transformation, which is under the control of a well-studied two-component quorum sensing signaling network (Claverys et al, 2009). Several lines of evidence have shown that the highly conserved ATP-dependent Clp protease, ClpP, in association with an ATPase subunit (either ClpC, ClpE, ClpL or ClpX), is involved in regulation of pneumococcal competence (Charpentier et al, 2000; Chastanet et al, 2001) (Fig. 6A). Identification of the ATPase subunit responsible for ClpP-dependent repression of competence was hampered because of the essentiality, depending on the growth medium and laboratory conditions, of several *clp* mutants including *clpP* and *clpX* (Chastanet et al, 2001). To address this issue, we employed CRISPRi and constructed sgRNAs targeting *clpP*, *clpC*, *clpE*, *clpL* and *clpX*. Competence development was quantified using a *luc* construct, driven by a competence-specific promoter (Slager et al, 2014). As shown in Fig. 6B, when expression of ClpP or ClpX was repressed by addition of IPTG, competence development was enhanced, while depleting any of the other ATPase subunits (ClpC, ClpE and ClpL) had no effect on competence (Fig. S15). This shows that ClpX is the main ATPase subunit responsible for ClpP-dependent repression of competence.

**Figure 6.**
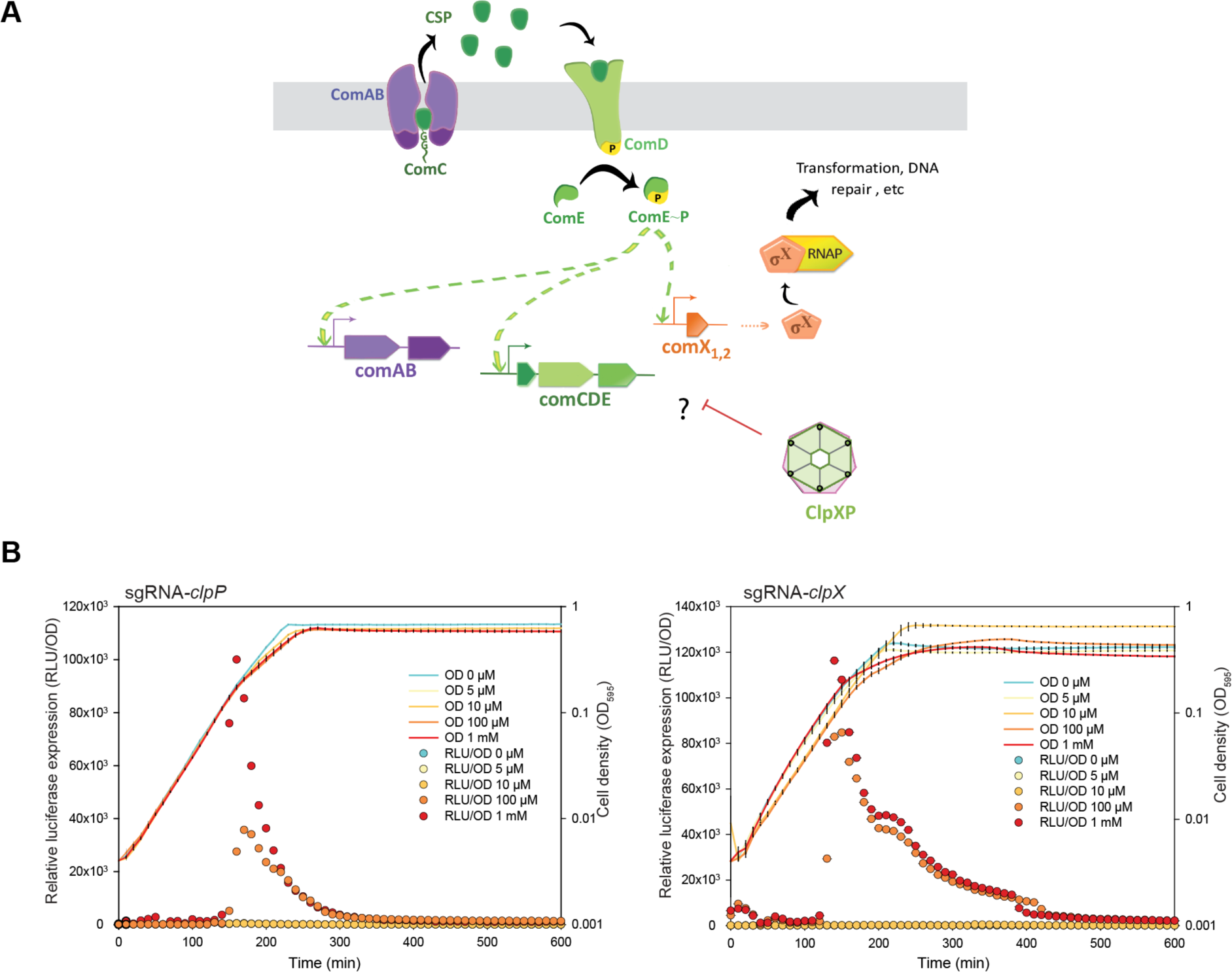
The ATPase ClpX and the ClpP protease repress competence development. **A**. Regulatory network of the competence pathway. Competence is induced when the *comC*-encoded competence-stimulating peptide (CSP), is recognized, cleaved and exported by the membrane transporter (ComAB). Accumulation of CSP then stimulates its receptor (membrane-bound histidine-kinase ComD), which subsequently activates ComE by phosphorylation, which in turn activates the expression of the so-called early competence genes. One of them, *comX*, codes for a sigma factor, which is responsible for the activation of over 100 late competence genes, including those required for transformation and DNA repair. Here, we show that the ATPase subunit ClpX, works together with the protease ClpP, repressing competence, probably by negatively controlling the basal protein level of the competence regulatory proteins but the exact mechanism is unknown (question mark). **B**. Repression of *clpP* or *clpX* by CRISPRi triggers competence development. Activation of competence system is reported by the *ssbB_luc* transcriptional fusion. Detection of competence development was performed in C+Y medium at a pH in which natural competence of the wild type strain is uninduced. IPTG was added to the medium at the beginning at different final concentrations (0, 5 ¼M, 10 ¼M, 100 ¼M, 1 mM). Cell density (OD595) and luciferase activity of the bacterial cultures were measured every 10 minutes. The value represents averages of three replicates with SEM.

### Concluding remarks

Here, we developed an IPTG-inducible CRISPRi system to study essential genes in *S. pneumoniae* (Fig. 1). In addition, we adopted a simple and efficient one-step sgRNA cloning strategy using infusion cloning, which results in ~89% positive sgRNA clones (Figs 2A and 2B), enabling high-throughput construction of sgRNA libraries. Using a stringent cut-off for phenotype detection, approximately 73% of the targeted genes showed growth phenotypes (Figs 2C, 2D and S2). It should be noted that polar effects of CRISPRi makes the phenotypes of genes in operons complicated to interpret. Also, after a delayed lag phase, most CRISPRi knockdowns with growth phenotypes eventually grow out to the same final OD indicating the emergence of a mutant subpopulation (Fig. S2B and S2C).

Based on analysis on the CRISPRi knockdowns, several previously ‘hypothetical’ genes could be functionally characterized and annotated (Figs S3-S10). For instance, combined with BlastP analysis and determination of *oriC-ter* ratios, we could annotate the pneumococcal primosomal machinery, including DnaA, DnaB, DnaC, DnaD, DnaG and DnaI (Table 1, Figs S3 and S13). Note that *spd2030* was mis-annotated as *dnaB*, which may be due to the different naming of primosomal proteins in *E. coli* and *Bacillus subtilis* (Briggs et al, 2012; Smits et al, 2011). By characterizing CRISPRi-based knockdowns with cell morphology defects, we identified 4 novel essential cell wall biosynthesis genes (*murT*, *gatD*, *tarP* and *tarQ*), which are promising candidates for future development of novel antimicrobials.

This work and other studies highlight that high-throughput phenotyping by CRISPRi is a powerful approach for hypothesis-forming and functional characterization of essential genes (Peters et al, 2016). We also show that CRISPRi can be used to unravel gene-regulatory networks in which essential genes play a part (Fig. 6). While we shed light on the function of just several previously uncharacterized essential genes, the here-described library contains richer information that needs to be further explored. In addition, CRISPRi screens can be used for mechanism of action (MOA) studies with new bioactive compounds. Indeed, CRISPRi was recently successfully employed to show that *B. subtilis* UppS is the molecular target of compound MAC-0170636 (Peters et al, 2016). We anticipate that the here-described pneumococcal CRISPRi library can function as a novel drug target discovery platform, can be applied to explore host-microbe interactions and will provide a useful tool to increase our knowledge concerning pneumococcal cell biology.

## Materials and Methods

### Strains, growth conditions and transformation

Oligonucleotides are shown in Supplementary Table 4 and strains in Supplementary Table 5. *S. pneumoniae* D39 and its derivatives were cultivated in C+Y medium, pH=6.8 (Slager et al, 2014) or Columbia agar with 2.5% sheep blood at 37ºC. Transformation of *S. pneumoniae* was performed as previously described (Martin et al, 2000), and CSP-1 was used to induce competence. Transformants were selected on Columbia agar supplemented with 2.5% sheep blood at appropriate concentrations of antibiotics (100 μg/ml spectinomycin, 250 μg/ml kanamycin, 1 μg/ml tetracycline, 40 μg/ml gentamycin, 0.05 μg/ml erythromycin). For construction of depletion strains with the Zn^2+^-inducible promoter, 0.1 mM ZnCl2/0.01 mM MnCl2 was added to induce the ectopic copy of the target gene. Working stock of the cells, called “T2 cells”, were prepared by growing the cells in C+Y medium to OD600 0.4, and then resuspending the cells with equal volume of fresh medium with 17% glycerol.

*E. coli* MC1061 was used for subcloning of plasmids and competent cells were prepared by CaCl2 treatment. The *E. coli* transformants were selected on LB agar with appropriate concentrations of antibiotics (100 μg/ml spectinomycin, 100 μg/ml ampicillin, 50 μg/ml kanamycin).

### Construction of an IPTG-inducible CRISPRi system in *S. pneumoniae*

*Streptococcus pyogenes dcas9* (*dcas9sp*) was obtained from Addgene (Addgene #44249, (Qi et al, 2013), and subcloned into plasmid pJWV102 (Veening laboratory collection) with the IPTG-inducible promoter PL*Sp* (Sorg, 2016) (here called PL) replacing PZn, resulting in plasmid pJWV102-PL-*dcas9sp*. pJWV102-PL-*dcas9sp* was integrated into the *bgaA* locus in *S. pneumoniae* D39 by transformation. To control PL expression, a codon-optimized *E. coli lacI* gene driven by the constitutive promoter F6 was inserted at the *prsA* locus in *S. pneumoniae* D39 (Sorg, 2016), leading to the construction of strain DCI23. DCI23 was used as the host strain for the insertion of gene-specific sgRNAs, and enables the CRISPRi system. The DNA fragment encoding the single-guide RNA targeting luciferase (sgRNA*luc*) was ordered as a synthetic DNA gBlock (Integrated DNA Technologies) containing the constitutive P3 promoter (Sorg et al, 2015). The sgRNA*luc* sequence is transcribed directly after the +1 of the promoter and contains 19 nucleotides in the base-pairing region, which binds to the non-template (NT) strand of the coding sequence of luciferase, followed by the optimized single-guide RNA used in Chen *et al.* 2013 (Chen et al, 2013), (Fig. S1A). Then, the sgRNA*luc* with P3 promoter was cloned into pPEP1 (Sorg et al, 2015) with removing the chloramphenicol resistance marker (pPEPX) leading to the production of plasmid pPEPX-P3-sgRNA*luc*, which integrates into the region between *amiF* and *treR* of *S. pneumoniae* D39. The pPEPX-P3-sgRNA*luc* is used as the template for generation of other sgRNAs by infusion cloning or inverse PCR method. The *lacI* gene with gentamycin resistance marker and flanked *prsA* regions was subcloned into pPEPY (Veening laboratory collection), resulting in plasmid pPEPY-PF6-*lacI.* This plasmid can be used to amplify *lacI* and integrate it at the *prsA* locus while selecting for gentamycin resistance. pJWV102-PL-*dcas9sp*, pPEPY-PF6-*lacI* and pPEPX-P3-sgRNA*luc* are available from Addgene (ID 85588, 85589 and 85590, respectively).

### Selection of essential genes

To identify each gene’s contribution to fitness for basal level growth, we performed Tn-Seq in *Streptococcus pneumoniae* D39 essentially as described before (Burghout et al, 2013; Zomer et al, 2012), but with growing cells in C+Y medium at 37ºC (data is available at SRA archive with accession number: SRP093201). Possibly essential genes were identified using ESSENTIALS (Zomer et al, 2012). We combined results of our Tn-Seq analysis in *S. pneumoniae* D39 with the results of two Tn-Seq studies in serotype 4 strain TIGR4 (van Opijnen et al, 2009; van Opijnen & Camilli, 2012), to select conserved essential genes of *Streptococcus pneumoniae* for CRISPRi. In the Tn-Seq study of 2012, fitness of each gene under 17 *in vitro* and 2 *in vivo* conditions was determined and genes were grouped into different classes (van Opijnen & Camilli, 2012). Based on that we defined a scoring system (essential as 1; likely essential as 0.75; core as 0.5; responsive as 0.3; responsive *in vivo* as 0.2; unresponsive as 0) and calculated the essentiality score for each gene (Supplementary Table 1). Finally, 391 genes were selected (Supplementary Table 1).

### Oligonucleotides for the CRISPRi library

*Design of sgRNAs*. The 20 nt guide sequences of the sgRNAs targeting different genes were selected with CRISPR Primer Designer developed by Yan et al. (Yan et al, 2015). Briefly, we searched within the coding sequence of each essential gene for a 14-nt specificity region consisting of the 12-nt ‘seed’ region of the sgRNA and GG of the 3-nt PAM (GGN). sgRNAs with more than one binding site within the pneumococcal genome, as determined by a BLAST search, were discarded. Next, we took a total length of 21-nt (including the +1 of the P3 promoter and 20-nt of perfect homology to the target) and the full-length sgRNA’s secondary structure was predicted using ViennaRNA (Lorenz et al, 2011), and the sgRNA sequence was accepted if the dCas9 handle structure was folded correctly (Larson et al, 2013). We chose the guide sequences as close as possible to the 5’ end of the coding sequence of the targeted gene (Qi et al, 2013). The sequences of the sgRNAs (20 nt) are listed in Supplemental Table S3.

### Cloning of sgRNA

We used infusion cloning instead of inverse PCR as recommended by Larson et al (Larson et al, 2013) because significantly higher cloning efficiencies were obtained with infusion cloning (SI text). Firstly, two primers, sgRNA_inF_plasmid_linearize_R and sgRNA_inF_plasmid_linearize_F, were designed for linearization of plasmid pPEPX-P3-sgRNA*luc*. These two primers bind directly upstream and downstream of the 19 bp guide sequence for *luc.* To fuse the 20-nt new guide sequence into the linearized vector, two 50-nt complementary primers were designed for each target gene. Each primer contains 15-nt at one end, overlapping with the sequence on the 5’ end of the linearized vector, followed by the 20-nt specific guide sequence for each target gene; and 15-nt overlapping with the sequence on the 3’ end of the linearized vector (Fig. 2B). The two 50-nt complementary primers were annealed in TEN buffer (10 mM Tris, 1 mM EDTA, 100 mM NaCl, pH 8) by heating at 95ºC for 5 min and cooling down to room temperature. The annealed product was fused with the linearized vector with the Quick-Fusion Cloning kit (Biotool, Cat. B22612) according to the manual, followed by direct transformation into *S. pneumoniae* D39 strain DCI23.

### Luciferase assay

*S. pneumoniae* strains XL28 and XL29 were grown to OD600=0.4 in 5 ml tubes at 37ºC, and then diluted 1:100 in fresh C+Y medium with or without 1 mM IPTG. Then, in triplicates, 250 μl diluted bacterial culture was mixed with 50 μl of 6×luciferin solution in C+Y medium (2.7 mg/ml, D-Luciferin sodium salt, SYNCHEM OHG) in 96-well plates (Polystyrol, white, flat, and clear bottom; Corning). Optical density at 595 nm (OD595) and luminescence signal were measured every 10 minutes for 10 hours using a Tecan Infinite F200 Pro microtiter plate reader.

### Growth assays

For growth curves of strains of the CRISPRi library (data for Figs 2C/D and S2B/C/D), T2 cells were thawed and diluted 1:1000 into fresh C+Y medium with or without 1 mM IPTG. Then 300 μl of bacterial culture was added into each well of 96-well plates. OD595 was measured every 10 minutes for 18 hours with a Tecan Infinite F200 Pro microtiter plate reader. For the data shown in Figs 3, S11 and S12, T2 cells were diluted 1:100 in C+Y medium. For growth assays of the depletion strains with the Zn^2+^ inducible promoter, T2 cells were thawed and diluted 1:100 into fresh C+Y medium with or without 0.1 mM ZnCl2/0.01 mM MnCl_2_.

### Detection of teichoic acids

#### Sample preparation

T2 cells of *S. pneumoniae* strains were inoculated into fresh C+Y medium with 0.1 mM Zn^2+^ by 1:50 dilution, and then grown to OD600 0.15 at 37ºC. Cells were collected at 8000 rcf for 3 min, and resuspended with an equal volume of fresh C+Y medium without Zn^2+^. Bacterial cultures were diluted 1:10 into C+Y with or without 0.1 mM Zn^2+^ or 1 mM IPTG (for CRISPRi strains), and then incubated at 37ºC. When OD600 reached 0.3, cells were centrifuged at 8000 rcf for 3 min. The pellets were washed once with cold TE buffer (10 mM Tris-Cl, pH 7.5; 1 mM EDTA, pH 8.0), and resuspended with 150 μl of TE buffer. Cells were lysed by sonication.

#### Detection of teichoic acid with phosphoryl choline antibody MAb TEPC-15

Protein concentration of the whole cell lysate was determined with the DC protein assay kit (Biorad Cat. 500-0111). Whole cell lysates were mixed with equal volumes of 2×SDS protein loading buffer (100 mM Tris-HCl, pH 6.8; 4% SDS; 0.2% bromphenolblue; 20% glycerol; 10 mM DTT), and boiled at 95°C for 5 min. 2 μg of protein were loaded, followed by SDS gel electrophoresis on a 12% polyacrylamide gel with cathode buffer (0.1M Tris, 0.1M Tricine, 0.1% SDS) on top of the wells and anode Buffer (0.2M Tris/Cl, pH9.9) in the bottom. After electrophoresis, samples in the gel were transferred onto a polyvinylidene difluoride (PVDF) membrane as described (Minnen et al, 2011). Teichoic acid was detected with anti-PC specific monoclonal antibody TEPC-15 (M1421, Sigma) by 1:1000 dilution as first antibody, and then with anti-mouse IgG HRP antibody (GE Healthcare UK Limited) with 1:5000 dilution as second antibody. The blots were developed with ECL prime western blotting detection reagent (GE Healthcare UK Limited) and the images were obtained with a Biorad imaging system.

### Microscopy

To detect the morphological changes after knockdown of the target genes, strains in the CRISPRi library were induced with IPTG and depletion strains were incubated in C+Y medium without Zn^2+^, stained with DAPI (DNA dye) and nile red (membrane dye), and then studied by fluorescence microscopy. Specifically, 10 μl of thawed T2 cells was added into 1 ml of fresh C+Y medium, with or without 1 mM IPTG, in a 1.5 ml Eppendorf tube, followed by 2.5 hours of incubation at 37ºC. After that, 1 μl of 1 mg/ml nile red was added into the tube and cells were stained for 4 min at room temperature. Then 1 μl of 1 mg/ml DAPI was added and the mix was incubated for one more minute. Cells were spun down at 8000 rcf for 2 min, and then the pellets were suspended with 30 μl of fresh C+Y medium. 0.5 μl of cell suspension was spotted onto a PBS agarose pad on microscope slides. DAPI, nile red and phase contrast images were acquired with a Deltavision Elite (GE Healthcare, USA). Microscopy images were analyzed with ImageJ.

For fluorescence microscopy of strains containing zinc-inducible GFP fusions, strains were grown in C+Y medium to OD600 0.1, followed by 10 time dilution in fresh C+Y medium with 0.1 mM Zn^2+^. After 1 hour of incubation, cells were spun down, washed with PBS and resuspended in 50 μl PBS. 0.5 μl of cell suspension was spotted onto a PBS agarose pad on microscope slides. Visualization of GFP was performed as described previously (Kjos et al, 2015).

For electron microscopy, T2 cells of *S. pneumoniae* strains were inoculated into C+Y medium with 0.1 mM Zn^2+^ and incubated at 37ºC. When OD600 reached 0.15, the bacterial culture was centrifuged at 8000 rcf for 2 min. The pellets were resuspended into C+Y without Zn^2+^ such that OD600 was 0.015, and then cells were incubated again at 37ºC. Bacterial cultures were put on ice to stop growth when OD600 reached 0.35. Cells were collected by centrifugation and washed once with distilled water. A small pellet of cells was cryo-fixed in liquid ethane using the sandwich plunge freezing method (Baba, 2008) and freeze-substituted in 1% osmium tetroxide, 0.5% uranyl acetate and 5% distilled water in acetone using the fast low-temperature dehydration and fixation method (McDonald & Webb, 2011). Cells were infiltrated overnight with Epon 812 (Serva, 21045) and polymerized at 60°C for 48 h. 90 nm thick sections were cut with a Reichert ultramicrotome and imaged with a Philips CM12 transmission electron microscope running at 90kV.

### Competence assays

The previously described *ssbB_luc* competence reporter system, amplified from strain MK134 (Slager et al, 2014), was transformed into the CRISPRi strains (sgRNA*clpP*, sgRNA*clpX,* sgRNA*clpL,* sgRNA*clpE,* sgRNA*clpC*, Supplementary Table 5). Luminescence assays for detection of activation of competence system were performed as previously described (Slager et al., 2014). IPTG was added into C+Y medium (pH 7.55) at the beginning of cultivation to different final concentrations.

## Acknowledgements

We thank A. Zomer, P. Burghout and P. Hermans (Radboud University, Nijmegen) for help with acquiring and analyzing the Tn-Seq data. We thank K. Kuipers and MI. de Jonge for providing the TEPC-15 antibodies and for the TA Western-blotting protocol. V. Benes and B. Haase (GeneCore, EMBL, Heidelberg) are thanked for sequencing support. XL is supported by China Scholarship Council (No. 201506210151). KK is supported by a NWO VENI fellowship (563.14.003). MK is supported by a grant from the research council of Norway (250976/F20). Work in the Veening lab is supported by the EMBO Young Investigator Program, a VIDI fellowship (864.12.001) from the Netherlands Organization for Scientific Research, Earth and Life Sciences (NWO-ALW), and ERC Starting Grant 337399-PneumoCell.

## Author Contributions

XL and JWV designed the study. XL, MK, ADP, CG, SPK, JS and JWV performed experiments and analyzed the data. KK performed electron microscopy. RAS developed the IPTG inducible system. JS analyzed the RNA-Seq data and performed the CRISPRi growth analysis. XL, JS, ADP, JRZ and JWV wrote the manuscript with input from all authors. All authors read and approved the final manuscript.

